# Effects of host plants nutrient on the nutrient in *Bradysia cellarum* and *Bradysia impatiens*

**DOI:** 10.1101/862094

**Authors:** Yuping Gou, Peter Quandahor, Yanxia Zhang, Changzhong Liu

## Abstract

The chive maggots *Bradysia cellarum* the fungus gnats *Bradysia impatiens* are two main root pests of plants. They can coexist on same host plants and have become devastating pests on liliaceous crops and edible fungi. Their growth and development are affected by nutrients of their host plants. We studied the effects of different host plant nutrients on the nutrient contents of these two *Bradysia* species. We assayed the nutrients in the roots of chive, board bean (B-bean), lettuce, cabbage, wild cabbage (W-cabbage) and pepper, and analysed the nutrient content of the two *Bradysia* species after three continuous generations of feeding on these different host plants. There chive and B-bean had higher contents of protein, free amino acid and starch than in other host plants. Soluble sugar, fat and protein contents were significantly higher in both *Bradysia* species when they were reared on chive and B-bean than when reared on cabbage, lettuce, W-cabbage and pepper. Our study provides a reference for further studies on the host range of the two *Bradysia* species, as well as knowledge for consideration in field crop rotations.

## Introduction

The chive maggots *Bradysia cellarum* Frey, 1948 (= *Bradysia odoriphaga* Yang and Zhang, 1985) and the fungus gnats *Bradysia impatiens* Johnnsen, 1912 (= *Bradysia difformis* Frey, 1948), are two main root insect pests which belong to the family Diptera and the genus Sciaridae [1, 2]. They may coexist on the same host plants and have become devastating pests on liliaceous crops and edible fungi [3]. The larvae of these species attack their host plants by chewing or stripping plant roots, especially the young and developing root hairs and stems of seedlings, and leading to losses in crop production and hindering agricultural productivity as well as farmers’ income [2, 4]. In outdoor fields, the two *Bradysia* species occur with similar regularities and outbreak in spring and autumn, while populations decline in summer [4]. The chive maggots *B. cellarum* heavily attacks chive, onion, garlic, cabbage and watermelon seedlings [5-8]. The fungus gnats *B. impatiens* [9] also causes damages to chive, lily, green onion, garlic, B-bean, cabbage, butterfly orchid and jonquil [10-13], which was first recorded on the edible mushroom in Yunnan, China in 2009 [14, 15].

Successful growth, development, and reproduction of plant-feeding insects depend on finding suitable host plants and obtaining adequate nutrition from them. It was found that there were significant differences among different host plants for the developmental duration, life longevity and population trend index of the fruit fly (*Bactrocera*), indicating that the fly’s population was influenced by host plant nutrition [16]. Protein, amino acid, soluble sugar and starch are the main nutrients in plants, which have a great influence on the growth, reproduction, survival rate and spawning of herbivores [17]. For example, higher soluble sugar content in wheat plants influenced a higher intrinsic rate of increase, shorter development time of the nymph and fecundity in the aphid species *Rhopalosiphum padi* L [18]. The soluble sugar content in the plant was negatively correlated with aphid resistance but positively correlated with fecundity [19, 20]. However, it was reported that the total carbohydrate contents of different plants had no significant effect on the growth and development of the beet armyworm *Spodoptera exigua* (Hubner), but was positively correlated with its oviposition and larval stage development [21]. Starch is a kind of polysaccharide in plant and plays an important role in storing energy. The starch content in a plant can be affected by insect attack, for instance, the starch content of wheat showed a decreasing trend after it was attacked by *Rhyzopertha dominica* (Fabricius) [22]. The nutrient content of host plants may also be one of the important factors that determines the host plant selection by insects [22].

Adults of *B. cellarum* and *B. impatiens* do not feed, so their main nutrients are accumulated and come from larval stages. Research on the biology [23], prevention and treatment [24], morphology [6] and sex pheromones [25] of *B. cellarum* and *B. impatiens* have attracted much more attention in recent years. However, the effects of host plant nutrients on the nutrient contents of the two *Bradysia* species have not been reported. Since different host plants have different nutrients and could differently affect the development of herbivorous insects, clear knowledge of the nutrient contents in different host plants and their correlations with insect feeding and growth is an important information for preventing and controlling insect pests of crops.

In the present study, we selected six host plants of *B. cellarum* and *B. impatiens*, including chive (*Allium tuberosum*), B-bean (*Vicia faba*), lettuce (*Lachca sativa*), cabbage (*Brassica pekinensis*), wild cabbage (*Brassica oleracea*) and pepper (*Capsicum annuum*), and determined the contents of protein, free amino acid, soluble sugar and starch in the roots of these six host plants. Additionally, we also measured these nutrient contents such as soluble sugar, glycogen, total fat, neutral fat and protein in these two *Bradysia* species after they were fed on these host plants for three continuous generations. The aim of this study was to explore the potential effects of host plant nutrients on the nutrient content of the two *Bradysia* species and to provide a reference for further studies on their host plant range, as well as knowledge for consideration in field crop rotations.

## Materials and methods

### Host plants

Seeds of chive (*Allium tuberosum*), B-bean (*Vicia faba*), lettuce (*Lachca sativa*), cabbage (*Brassica pekinensis*), W-cabbage (*Brassica oleracea*) and pepper (*Capsicum annuum*) were sown in the laboratory of Gansu Agricultural University (36°5’20” N, 103°41’54” E), Lanzhou, China. Each seeds planted three pots. Two months after germination, the stems of each host plants were collected to feed larvae of the two *Bradysia* species. Plant roots were extracted and cleaned with water. Their nutrient contents were then determined individually.

### Insects

Populations of *B. cellarum* and *B. impatiens* were collected from chive fields in Gangu county (34°44’44” N, 105°20’13” E), Gansu Province, China. Larvae were reared with stems of each tested host plants in moisturized petri dishes [26-27] for 3 continuous generations before being used in experiments.

### Determination of nutrients in tested plants

The roots from three pots of the six host plants were used to determine the contents of soluble protein, free amino acid, soluble sugar and starch separately. Each experiment was repeated 3 times. The protein content was measured using the Bradford assay method [28]. The optical density (OD) values were measured at 595 nm, and the protein contents were calculated based on the standard curve of BSA (Sangon biotech, Shanghai, China). The free amino acid content was detected by the ninhydrin method [28]. The OD value was measured at 580 nm, and calculated based on the standard curve of leucine. The soluble sugar content was determined using the anthrone colorimetry method and calculated based on the standard curve of D-glucose (Sangon biotech, Shanghai, China) [28].

### Carbohydrate determination in the insect body

The soluble sugar and the glycogen contents of the two *Bradysia* species reared with different host plants were determined using the methods of Lv et al [29]. Firstly, we transferred 160 µL of the homogenate suspension into a 2 mL centrifuge tube and mixed it with 20 µL of 20% sodium sulphate and 1500 µL of a chloroform/methanol solution (1:2 v/v). Then, this mixture was centrifuged at 10,000 rpm for 15 min at 4 °C. Hereafter, this top mixture solution was treated as a premeasured solution.

In order to determine the soluble sugars of *Bradysia* larvae, firstly, we transferred 150 µL suspension of the premeasured solution into a 1.5 mL centrifuge tube and then mixed it with 10 µL ddH_2_O and 240 µL anthrone (1.42 g/L). After which the mixture was first incubated for 10 min at 25 °C and then incubated in boiling water for another 10 min, followed by cooling to room temperature. The mixture was then transferred into a 96-pore coated plate. Finally, the OD value was determined at 620 nm wavelength, and the soluble sugar content was calculated based on the standard curve of D-glucose (Sangon biotech, Shanghai, China).

To determine the glycogen of *Bradysia* larvae, similarly, the remaining suspension of the premeasured solution was transferred to another centrifuge tube. The precipitant was mixed with 400 µL of 80% methanol and homogenized in an ultrasonic cleaning apparatus for 5-10 min, after which the homogenate was centrifuged again at 10,000 rpm for 10 min at 4 °C. Then, the new suspension was mixed with anthrone solution, incubated for 10 min under room temperature, and then in boiling water for another 10 min. the mixture was then transferred into a 96-pore coated plate. The OD value was measured at 620 nm, and the glycogen content was calculated based on the standard curve of D-glucose. Each experiment was repeated three times and each time with fifteen third instar larvae of *B. cellarum* and *B. impatiens*.

### Total fat and neutral fat determination in the insect body

The total fat and neutral fat of *Bradysia* larvae after feeding on different host plants were determined using the methods of Lv et al [29]. To determine total fat content, 100 µL of the premeasured solution was transferred to a 1mL centrifuge tube, then dried at 90 °C until all the solvents were completely evaporated. After that, 10 µL of 98% sulphuric acid and 190 µL of vanillin solution (1.2 g/L) were added to the tube. After 15 min of incubation under room temperature, the liquid mixture was transferred into a 96-pore coated plate. The OD value of total fat content was determined at 525 nm and calculated based on the standard curve of triolein (Sigma, St. Louis, MO, USA). The experiment was repeated three times and each time with fifteen third instar *Bradysia* larvae.

For the detection of neutral fat in the larvae of the two *Bradysia* species, firstly, approximately 150 µL of the premeasured solution was transferred into a 1.5 mL centrifuge tube and then dried at 90 °C until all the solvents were completely evaporated. Following this, 1 mL of chloroform was added to dissolve the solution content and then centrifuged at 10,000 rpm for 15 min at 4 °C. After which 100 µL of the new suspension was treated in the same procedure as for total fat detection. The OD value of neutral fat was measured at 525 nm, and its content was calculated based on the standard curve of triethylhexanoin (Aladdin, Shanghai, China). The experiment was repeated three times and each time with fifteen 3th instar *Bradysia* larvae.

### Protein determination in insect body

The protein content of the two *Bradysia* species in the different treatments was determined referring to the method of Lv et al [29]. About 20 µL of the homogenate suspension was transferred into a 96-well plate and then mixed with 200 µL of Coomassie brilliant blue G-250dye (Bradford assay) for 15-20 min. The OD values were measured at 595 nm, and the protein contents were calculated based on the standard curve of BSA (Sangon biotech, Shanghai, China). The experiment was repeated three times with fifteen third instar larvae of *Bradysia* each time.

### Statistical analysis

Microsoft Excel 2016 was used to analyse data and plot figures. Proc Means program (SPSS 19.0, IBM, Armonk, NY, USA) was used for one-way analysis of variance [30] and Tukey’S HSD was used to compare the differences among variables [31].

## Results

### Nutrient contents in roots of different host plants

The root protein content in the 6 tested host plants are shown in Figure 1A. The B-bean had the highest content (4.49 mg/g), followed by chive (3.87 mg/g), pepper (2.34 mg/g), W-cabbage (1.99 mg/g), lettuce (1.91 mg/g) and cabbage (1.61 mg/g). The protein contents in B-bean and chive were significantly (P< 0.05) higher than those of the other 4 host plants (Fig 1A).

**Fig 1.**
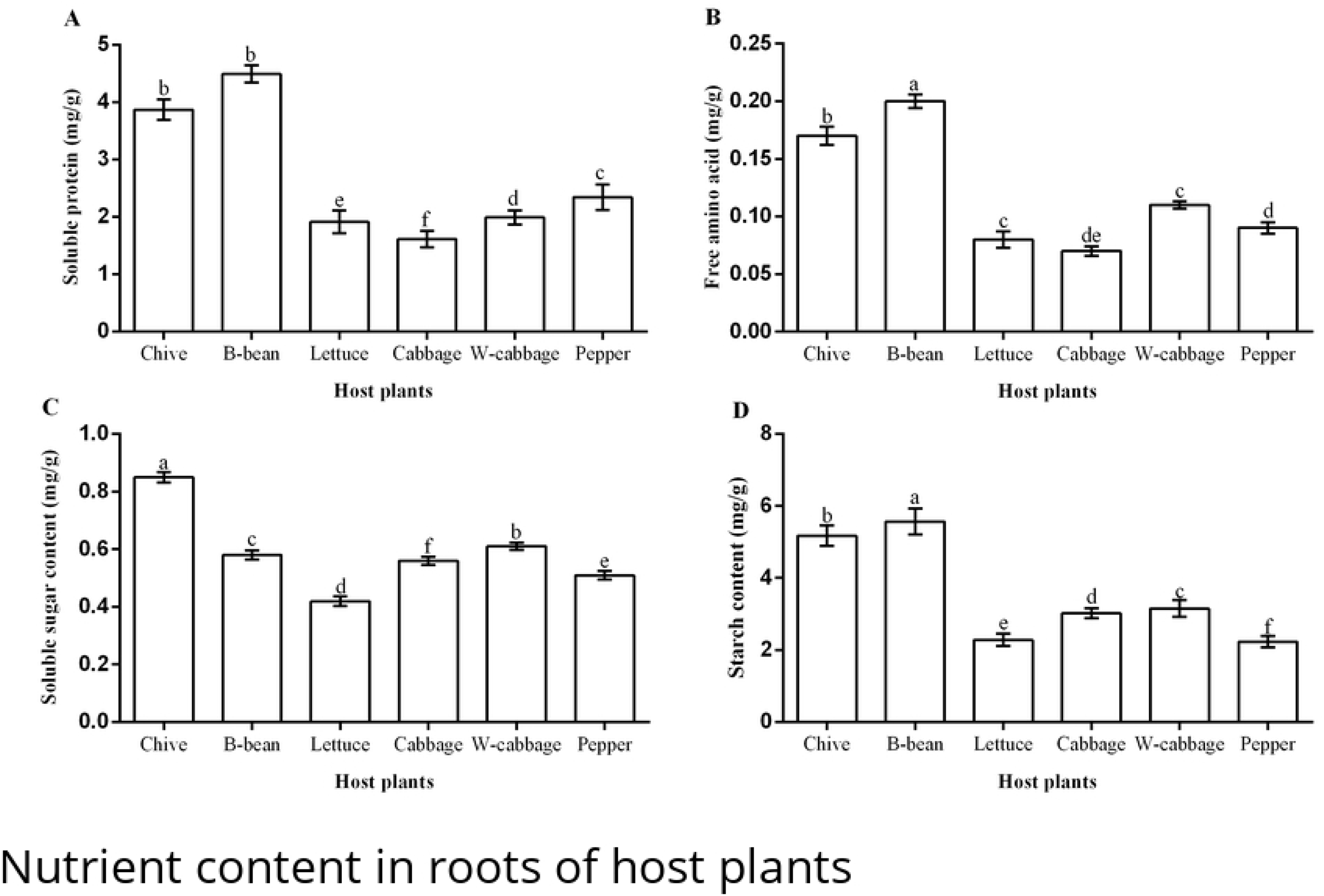
Nutrient contents in roots of host plant. A: Protein content; B: Free amino acid content; C: Soluble sugar content; D: Starch content. Different small letters indicate significant difference at the 0.05 level among treatments of different host plants (Tukey’s HSD).

Free amino acid contents in the order from high to low were 0.20 mg/g in B-bean, 0.17 mg/g in chive, 0.11 mg/g in W-cabbage, 0.09 mg/g in pepper, 0.08 mg/g in lettuce and 0.07mg/g in cabbage (Fig 1B). The roots soluble sugar content in the 6 tested host plants were 0.85 mg/g in chive, followed by 0.61 mg/g in W-cabbage, 0.58 mg/g in B-bean, 0.56 mg/g in cabbage, 0.51 mg/g in pepper and 0.42 mg/g in lettuce (Fig 1C). The highest starch content among the tested host plants occurred in the roots of B-bean (5.57 mg/g), while the lowest starch content occurred in the roots of pepper (2.23 mg/g) (Fig 1D). The roots starch content of other host plants were 5.18 mg/g of chive, 3.15 mg/g of W-cabbage, 3.02 mg/g of cabbage and 2.28 mg/g of lettuce. The starch content was significantly (P<0.05) higher in the roots of B-bean and chive than those of other 4 host plants.

### Soluble sugar contents in *B. cellarum* and *B. impatiens*

Soluble sugar contents of *B. cellarum* and *B. impatiens* reared on the six host plants are showed in Fig 2. We found that soluble sugar content of *B. cellarum* reared on chive was the highest (8.76 µg/mg). This was followed by 7.57 µg/mg, 7.53 µg/mg and 6.38 µg/mg if they were reared on lettuce, B-bean and W-cabbage, respectively. The soluble sugar contents were even lower when reared on pepper (5.91 µg/mg) and cabbage (5.78 µg/mg). However, the highest soluble sugar content of *B. impatiens* occurred on B-bean (9.13 µg/mg), followed by chive (8.95 µg/mg), lettuce (7.97 µg/mg), W-cabbage (7.34 µg/mg) and cabbage (5.68 µg/mg). Moreover, the soluble sugar content of *B. impatiens* was the lowest when reared on pepper (4.93 µg/mg). There were significant differences in the soluble sugar contents between *B. cellarum* and *B. impatiens* when fed on B-bean and pepper (P< 0.05). We also found that *B. cellarum* and *B. impatiens* obtained higher soluble sugar contents when fed on chive, B-bean and lettuce compared with when fed on cabbage, W-cabbage, and pepper.

**Fig 2.**
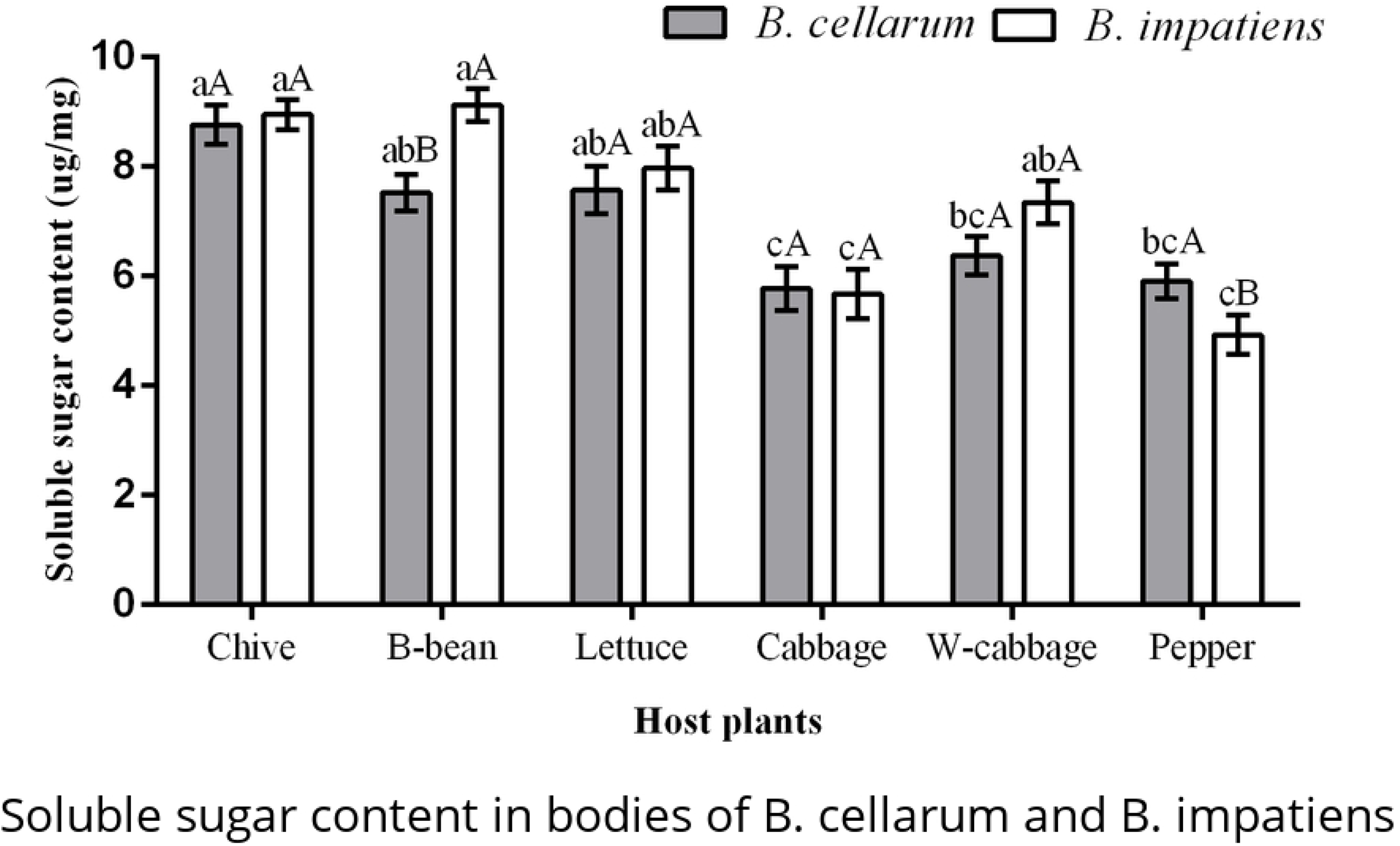
Soluble sugar content in bodies of *B. cellarum* and *B. impatiens*. Values are the means ± standard error. Different lowercase letters indicate significant differences between two *Bradysia* species on different host plants by Tukey’s HSD (P< 0.05); while the different uppercase letters represent significant differences between two *Bradysia* species on same host plants by Tukey’s HSD (P < 0.05).

### Glycogen contents in *B. cellarum* and *B. impatiens*

The glycogen contents of *B. cellarum* and *B. impatiens* across the various host plants are presented as Fig 3. The glycogen content of *B. cellarum* was the highest reared on lettuce (6.37 µg/mg), followed by board bean (5.36 µg/mg), chive (4.44 µg/mg), W-cabbage (4.43 µg/mg), and pepper (3.83 µg/mg), while the lowest occurred on cabbage (3.34 µg/mg). Glycogen contents of *B. impatiens* reared on the six host plants in order from the highest to the lowest were 6.50 µg/mg on B-bean, 6.106 µg/mg on lettuce, 4.28 µg/mg on chive, 4.26 µg/mg on W-cabbage, 3.42 µg/mg on pepper and 2.86 µg/mg on cabbage. Meanwhile, significant (P< 0.05) differences in the glycogen content of *B. cellarum* were observed when reared on lettuce, chive, cabbage, W-cabbage and pepper, and in the glycogen content of *B. impatiens* fed on chive, B-bean, cabbage, W-cabbage and pepper. In addition, there were significant (P< 0.05) differences in the glycogen contents between *B. cellarum* and *B. impatiens* when they were fed on B-bean and pepper. Moreover, we found that *B. cellarum* and *B. impatiens* obtained higher glycogen when they were fed on chive, B-bean, lettuce and W-cabbage than when fed on cabbage and pepper.

**Fig 3.**
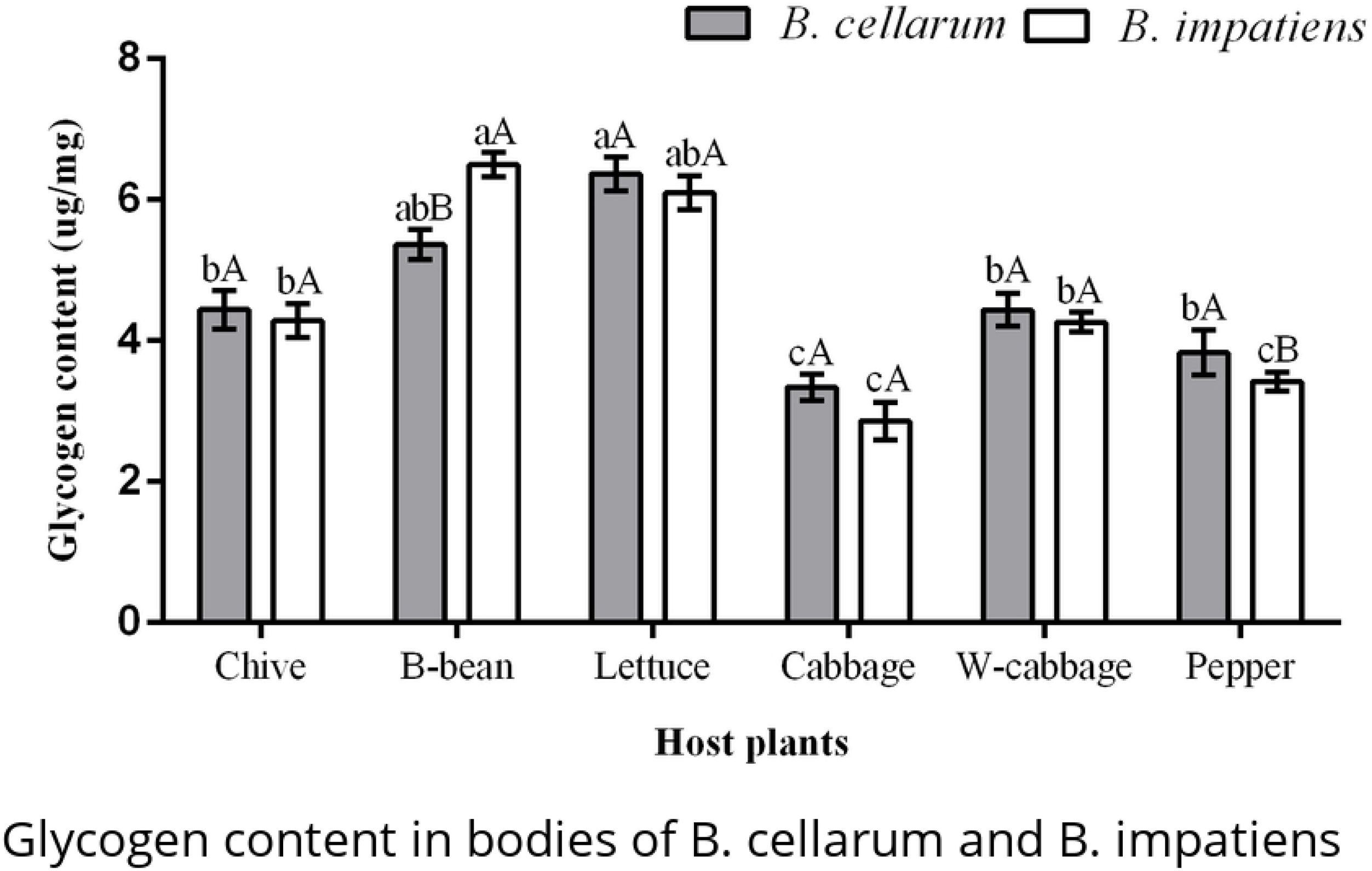
Glycogen content in bodies of *B. cellarum* and *B. impatiens*. Values are the means ± standard error. Different lowercase letters indicate significant differences between two *Bradysia* species on different host plants by Tukey’s HSD (P< 0.05); while the different uppercase letters represent significant differences between two *Bradysia* species on same host plants by Tukey’s HSD (P < 0.05).

### Total fat contents in *B. cellarum* and *B. impatiens*

Total fat contents of *B. cellarum* and *B. impatiens* reared on six host plants are showed in Fig 4. The highest total fat content of *B. cellarum* was 8.43 µg/mg when fed on chive. Those reared on B-bean, cabbage, pepper and W-cabbage exhibited lower total fat content of *B. cellarum* of 7.67 µg/mg, 7.00 µg/mg, 6.66 µg/mg and 6.42 µg/mg, respectively. Total fat content of *B. cellarum* was the lowest (5.56 µg/mg) when fed on lettuce. Total fat content of *B. impatiens* reared on chive was 8.64 µg/mg, which was higher in comparison with those of *B. impatiens* fed on B-bean (7.83 µg/mg), pepper (7.24 µg/mg), W-cabbage (6.99 µg/mg) and cabbage (6.34 µg/mg). Like *B. cellarum, B. impatiens* also had the lowest total fat content when reared on lettuce (5.97 µg/mg). In terms of the total fat contents of *B. cellarum* and *B.impatiens* reared on the same host plants, significant (P< 0.05) differences were observed across cabbage, W-cabbage and pepper.

**Fig 4.**
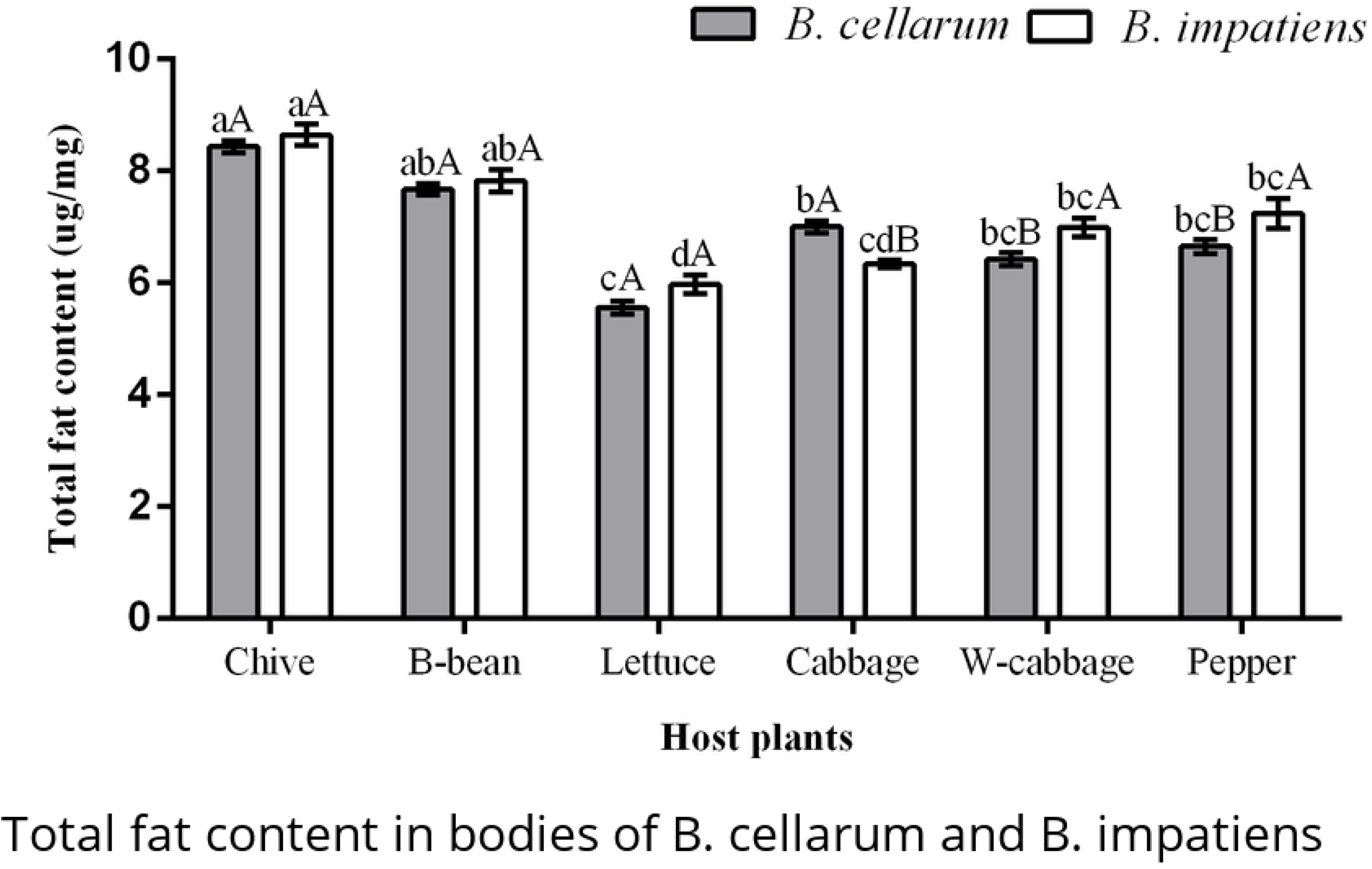
Total fat content in bodies of *B. cellarum* and *B. impatiens*. Values are the means ± standard error. Different lowercase letters indicate significant differences between two *Bradysia* species on different host plants by Tukey’s HSD (P< 0.05); while the different uppercase letters represent significant differences between two *Bradysia* species on same host plants by Tukey’s HSD (P < 0.05).

### Neutral fat contents in *B. cellarum* and *B. impatiens*

Feeding on six host plants exerted influences on the neutral fat contents of *B. cellarum* and *B. impatiens* (Fig 5). We found that neutral fat content of *B. cellarum* was significantly higher when reared on chive (0.92 µg/mg) and B-bean (0.91 µg/mg) than on pepper (0.74 µg/mg) and cabbage (0.69 µg/mg). Neutral fat content of *B. cellarum* reared on lettuce was the least (0.52 µg/mg), followed by the W-cabbage (0.55 µg/mg). Similar to *B. cellarum*, the neutral fat content of *B. impatiens* was higher when reared on chive and B-bean, with 0.95 and 0.84 µg/mg, respectively, which significantly differed from when fed on pepper (0.73 µg/mg), cabbage (0.64 µg/mg), W-cabbage (0.53 µg/mg) and lettuce (0.50 µg/mg). For the neutral fat contents of *B. cellarum* and *B. impatiens* reared on same host plants, there was significant (P< 0.05) difference between two species only when fed on B-bean. There was no significant difference when fed on chive, lettuce, cabbage, W-cabbage and pepper.

**Fig 5.**
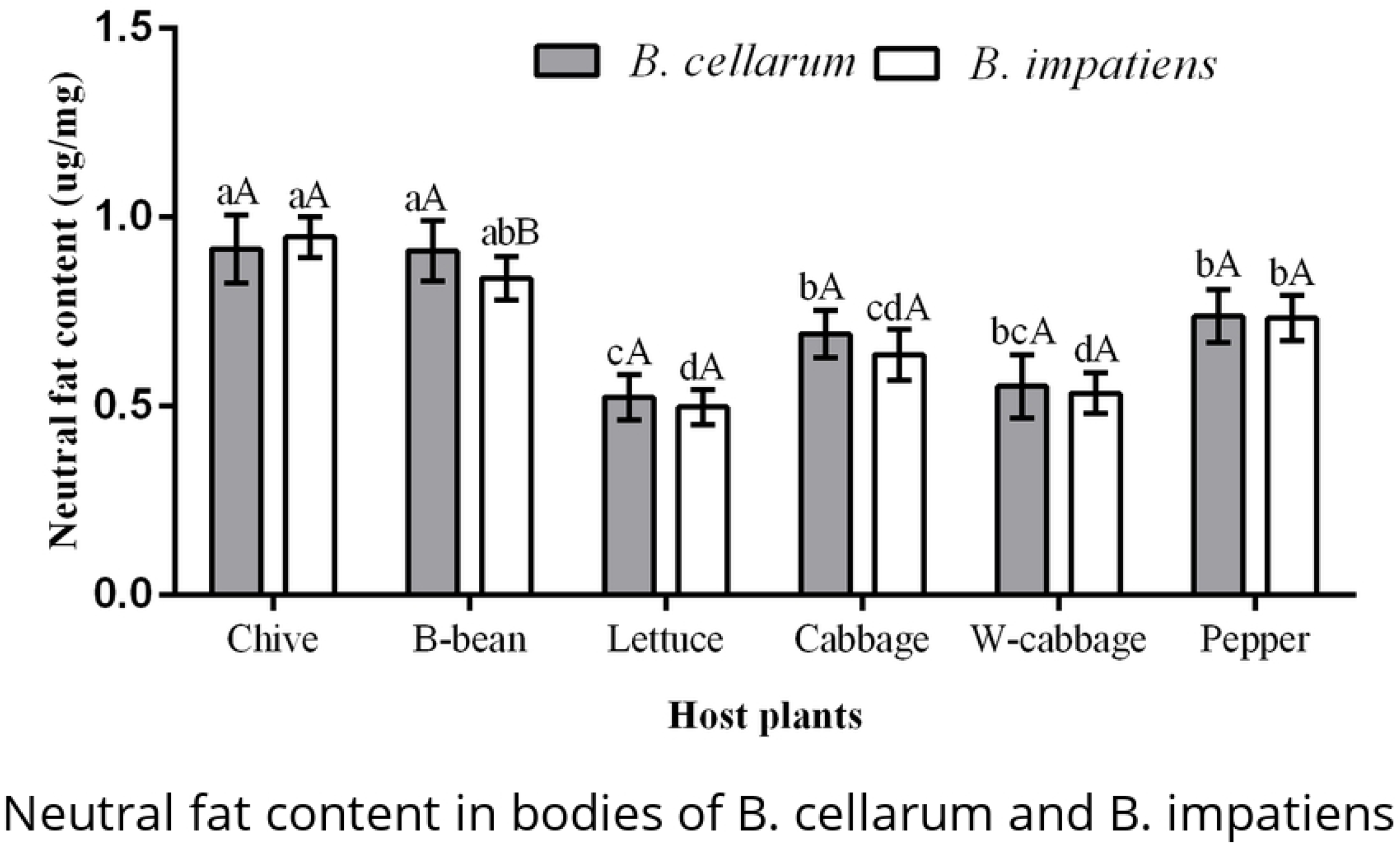
Neutral fat content in bodies of *B. cellarum* and *B. impatiens*. Values are the mean ± standard error. Different lowercase letters indicate significant differences between two *Bradysia* species on different host plants by Tukey’s HSD (P< 0.05); while the different uppercase letters represent significant differences between two *Bradysia* species on same host plants by Tukey’s HSD (P < 0.05).

### Protein contents in *B. cellarum* and *B. impatiens*

The protein contents of *B. cellarum* were much higher when reared on B-bean (43.94 µg/mg) and chive (43.57 µg/mg), and significantly (P< 0.05) differed from when reared on pepper (34.67 µg/mg), lettuce (34.47 µg/mg), cabbage (29.79 µg/mg) and W-cabbage (28.99 µg/mg) (Fig 6). The protein content of *B. impatiens* was the highest when fed on B-bean (51.61 µg/mg) and remarkably (P< 0.05) differed from when reared on chive (43.02 µg/mg), W-cabbage (37.17 µg/mg), cabbage (37.02 µg/mg), lettuce (36.99 µg/mg) and pepper (29.69 µg/mg). In addition, we also found that the protein contents of *B. cellarum* and *B. impatiens* reared on the same host plants showed significant difference between two species occurred on cabbage and W-cabbage. There was no significant differences when they were fed on chive, B-bean, lettuce and pepper.

**Fig 6.**
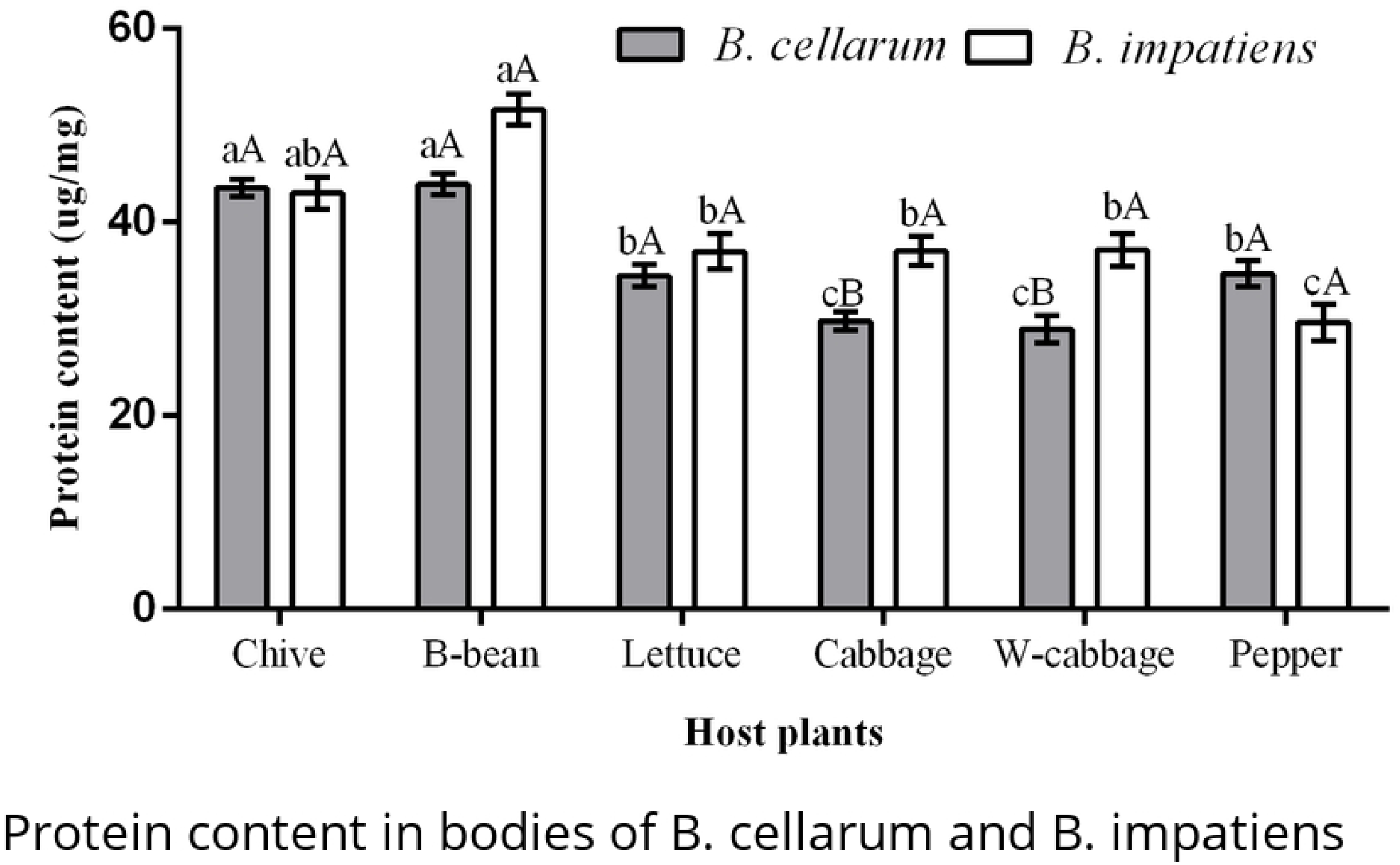
Protein content in bodies of *B. cellarum* and *B. impatiens*. Values are the mean ± standard error. Different lowercase letters indicate significant differences between two *Bradysia* species on different host plants by Tukey’s HSD (P< 0.05); while the different uppercase letters represent significant differences between two *Bradysia* species on same host plants by Tukey’s HSD (P < 0.05).

### Correlation analysis between host plant nutrients and body contents in *B. cellarum* and *B. impatiens*

The protein contents in the body of *B. cellarum* were significantly positively correlated with the protein contents (P< 0.01) and the free amino acid content (P< 0.05) of the host plant roots, with correlation coefficients of 1.00 and 0.868, respectively (Table 1). The neutral fat content of *B. cellarum* body had a significantly (P<0.05) positive correlation with the free amino acid contents and the starch content of the host plant roots with correlation coefficients of 0.840 and 0.823 respectively (Table 1). The total fat contents of *B. cellarum* body were significantly correlated with the contents of soluble sugars and starch. The glycogen content of *B. cellarum* body had a negative correlation with plant soluble sugar content. The starch content in the host plant roots was significantly correlated with the content of total fat and neutral fat in the insect body.

**Table 1.**
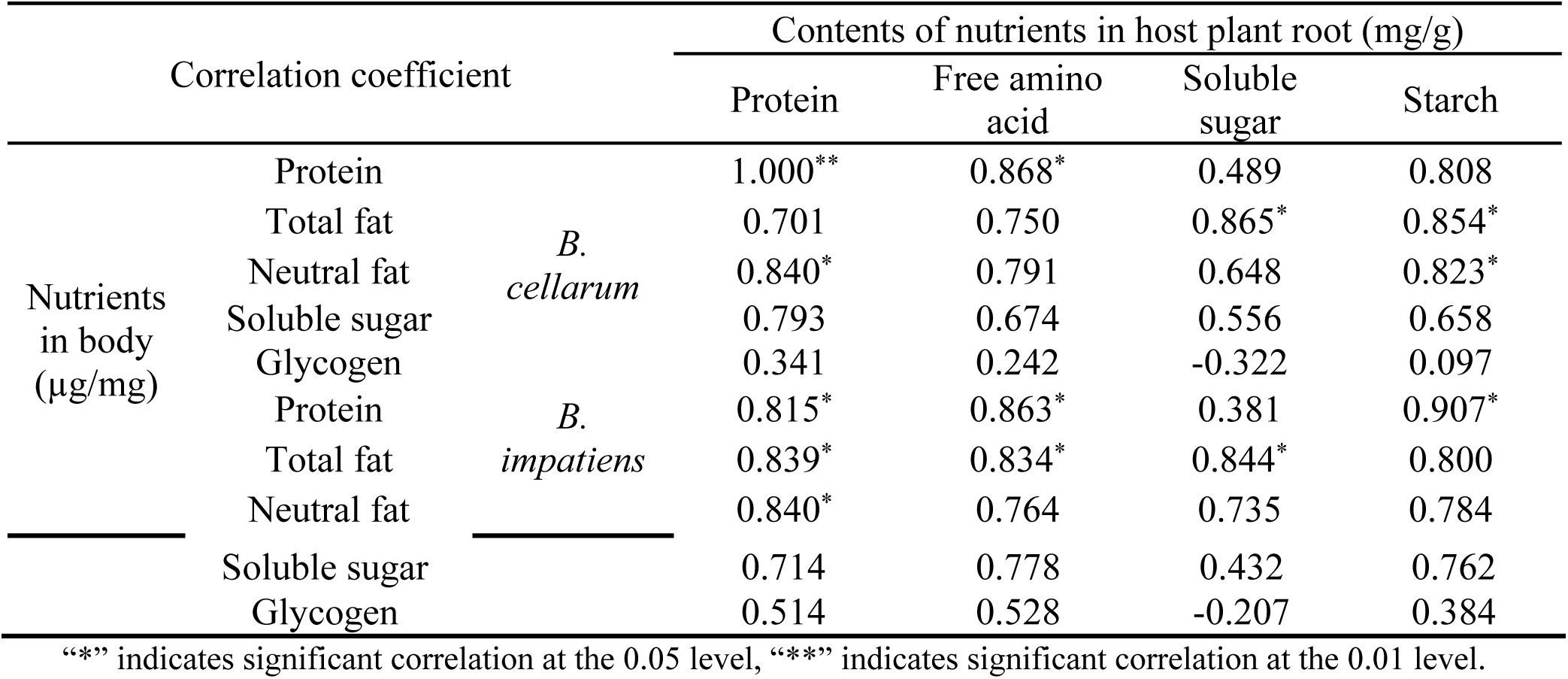
Correlation analysis between nutrient in host plant roots and nutrient in *B. cellarum* and *B. impatiens*.

The contents of protein, total fat and neutral fat in the body of *B. impatiens* were all significantly positively correlated with the protein content of the host plant roots with correlation coefficients of 0.815, 0.839 and 0.840, respectively (Table 1). The contents of protein and total fat in the *B. impatiens* body was significantly (P< 0.05) positively correlated with the free amino acid content of host plant roots with correlation coefficients of 0.863 and 0.834 respectively. The contents of protein and total fat in the body of *B. impatiens* were significantly (P< 0.05) positively correlated with the soluble sugar in host plant roots. The glycogen content of *B. impatiens* body was negatively correlated with the soluble sugar in host plant roots. There was an extremely significant (P< 0.01) correlation between the protein content of *B. impatiens* and the starch content in roots with a correlation coefficient of 0.907.

## Discussion

In this study, we found that contents of protein and free amino acid were much higher in B-bean and chive, compared to the other four host plants. Our findings also revealed that the two *Bradysia* species obtained much more protein contents when they were reared on chive and B-bean. Furthermore, we found that the contents of protein and free amino acid in plant roots were significantly correlated with the protein contents in the body of two *Bradysia* species. It has been reported that *B. cellarum* and *B. impatiens* had a stronger adaptability with shorter developmental duration and higher oviposition, net reproductive rate and intrinsic rate value when they fed on chive and B-bean [11]. This probably could be a result of the high level of protein and free amino acid in these host plants which promote the growth, development and fecundity of the *Bradysia* species. It was reported that the western flower thrip *Frankliniella occidentalis* moved to pollen from leaves once the plants bloomed because the pollen contained higher protein than leaves [32]. Therefore, we presume that the two *Bradysia* species prefer a host plant with high protein and free amino acid contents.

Our results revealed that the soluble sugar contents were significantly higher in chive, W-cabbage and B-bean than lettuce, cabbage and pepper. Moreover, we also found that the soluble sugar content in the body of the two *Bradysia* species especially *B. impatiens* was the highest when they were reared on the chive and B-bean. This is consistent with the report that *B. impatiens* heavily attacked chive and B-bean [13]. Other studies revealed that the soluble sugar content in crop was negatively correlated with aphid resistance. That is, the crop with higher soluble sugar content were less resistant to aphids. This was because the amount of soluble sugar positively promoted the fecundity of aphids [21, 33, 34]. A recent study reported that *B. cellarum* reared on chive had significantly longer female and male longevity, as well as higher oviposition and survival rate [35]. However, our correlation analysis showed that the soluble sugars may have contributed mainly to the total fat content in the *Bradysia* species. This is consistent with the report that carbohydrate provides energy for insect growth and flight [21]. However, an interesting finding from the current study revealed that the soluble sugar content in the host plant roots were inversely correlated with the glycogen in the body of both *Bradysia* species. Glycogen is the energy storage material of insects, known as animal starch, which is used as the energy storage material for embryo development, and is correlated with the reproduction of insects [36]. It is also the essential animal version of starch and the main energy storage material in organisms [36]. There may be a trade-off between the nutrient requirement and the reproduction of the two *Bradysia* species, as the adults do not feed.

Our results show that the two *Bradysia* species select host plants for their higher soluble sugar content and protein content. The larvae of *B. cellarum* reared on chive had much higher soluble sugar content than those reared on B-bean. However, the larvae of *B. impatiens* reared on B-bean had much higher soluble sugar content than those reared on chive. The results also showed that *B. cellarum* obtained a higher glycogen content when they were reared on lettuce. While *B. impatiens* obtained it when they were reared on B-bean. These physiological differences between two *Bradysia* species is worthy of further investigation.

In summary, the nutrient content in an insect body is an important guarantee for their growth, development and reproduction. Our study demonstrated that there are strong correlations between host plant nutrients, especially proteins and soluble sugars, with the nutrient content in *Bradysia* species. The host plants nutrients, especially soluble sugar and soluble protein, may have strongly affected the nutrient contents of *B. cellarum* and *B. impatiens.* Since the contents of protein and free amino acid are much higher in chive and B-bean, it is proposed that rotation of those two plants should be avoided in the field to control for rapid population expansion and the damage caused by *B. cellarum* and *B. impatiens*. Such strategy could cause a decrease in the continuous supply of their nutrient requirements from these host plants, which could further weaken their behaviour and performance. This approach could be a novel strategy for pest control.

## Acknowledgments

We acknowledge Prof Jing-Jiang Zhou of Rothamsted Research, UK for critical reading and helpful suggestions of this article. We are also grateful to all editors and reviewers for handling our manuscript. We appreciate the proofreading by entomology editing.

